# Root interactions and plant growth in a tomato/potato onion intercropping system under different phosphorus levels

**DOI:** 10.1101/142067

**Authors:** Hongjie Yu, Shaocan Chen, Xingang Zhou, Fengzhi Wu

## Abstract

Intercropping systems have been implemented in many parts of the world due to their beneficial effects on yield and biomass. In intercropping systems, changes in plant growth are usually related to variations in root distribution and phosphorus (P) levels, however, root distributions and root tendencies are difficult to study, as root systems grow beneath the soil surface. Therefore, we have a relatively poor understanding of the relationship between plant root interactions and plant growth in intercropping systems. In this study, a custom apparatus consisting of a transparent manual root box was used to observe intact root systems in situ. We investigated how root distribution and root tendency changed in a tomato/potato onion intercropping system under two P treatments, and how tomato plant growth was influenced. The results showed that the shoot and total biomass for the tomato plants were increased by intercropping with potato onion plants under both tested P levels, the root distribution of intercropped tomato plants was deeper than that of monocultured plants, and the tomato roots tended to grow away from the potato onion roots. Our research reveals that a deeper and more evasive root distribution is more conductive to the plant growth of the intercropped tomato.

**SUMMARY STATEMENT:** Our study can help more people clearly know the relationship between the root interactions and plant growth in mixed cultures.

## INTRODUCTION

Roots are extremely important plant organs that mediate nutrient and water absorption (Barber, 1995; Fang et al., 2011). Over the long evolutionary history of plants, different species have developed varying degrees of root plasticity. Within the complex living environment, root plasticity can be influenced by many factors (Karban, 2008), such as nutrient heterogeneity (Fang et al., 2009; Hodge, 2004; Liao et al., 2001, 2004) and the presence of neighboring roots (de Kroon et al., 2003; Dudley and File, 2007; Falik et al., 2003; Karban and Shiojiri, 2009; Maina et al., 2002).

In recent years, the topic of root recognition has attracted the attention of a growing number of scholars. The presence of a neighboring plant that represents a resource competitor can trigger an increase in root biomass allocation (Falik et al., 2003; Gersani et al., 2001; Maina et al., 2002; O’Brien et al., 2005; Padilla et al., 2013), whereas some plants can recognize other individuals of their own species and limit root proliferation (Biedrzycki and Bais, 2010; Biedrzycki et al., 2010; Dudley and File, 2007). For example, when planted with non-kin species instead of siblings, the Great Lakes Sea Rocket (*Cakile edentula*) accumulates more biomass in its fine roots (Dudley and File, 2007). In addition, *Impatiens pallida* plants are capable of kin recognition only when the roots of another plant are present (Murphy and Dudley, 2009). Therefore, when analyzing intercropping systems, which are practiced in many parts of the world, including tropical, subtropical and temperate regions (Francis, 1986; Vandermeer, 1989), the process of root recognition should be considered.

Intercropping can confer considerable yield and biomass advantages in certain situations (Awal et al., 2006; Tsubo and Walker, 2002; Zhang et al., 2007). With respect to intercropping systems, most studies have addressed interspecific facilitation, in which different plant species benefit one another when two species are grown together. Interspecific facilitation can benefit plant growth and nutrient absorption after intercropping with other species in agro-ecosystems (Ae et al., 1990; Cu et al., 2005; Gardner and Boundy, 1983; He et al., 2013; Horst and Waschkies, 1987; Li et al., 2001, 2003, 2007, 2010). However, as root systems are hidden below ground, it can be difficult to observe and quantify root growth in situ, explaining why there are relatively few studies concerning root interactions in these important agricultural systems. Li et al. (2006) and Gao et al. (2010) confirmed that greater lateral root deployment and compatibility of spatial root distribution in intercropping species contribute to higher yields and plant growth. Xia et al. (2013) suggested that total root length growth and the spatial distribution of roots are sensitive to phosphorus (P) application in cropping systems (Xia et al., 2013). Phosphate ions in soil usually become unavailable by reacting with soil cations to form either soluble complexes or insoluble precipitates (Cu et al., 2005), or they adsorb to the surfaces of various positively charged soil particles (Hinsinger et al., 2003). Based on these investigations, we can conclude that plant root behavior is much more complex than previously thought.

Tomato (*Solanum lycopersicum* L.) is a widely cultivated vegetable around the world, although continuous monocropping of tomato plants and excessive fertilizer application have resulted in soil acidification and salinization in many locations, decreasing tomato yields and fruit quality (Liu et al., 2014). The potato onion (*Allium cepa* L. var. *aggregatum* G. Don) is an onion variety that is widely cultivated in northeastern China and is a good companion plant for tomato. In many previous studies, intercropping of tomatoes and potato onions has been shown to increase tomato quality, alleviate tomato *Verticillium* wilt and improve soil quality by altering soil enzyme activities and microbial communities (Fu et al., 2015, 2016; Liu et al., 2014; Tringovska et al., 2015; Wu et al., 2013, 2016). However, we know little about how the roots of these species interact when tomato plants are intercropped with potato onion plants, and in particular, it is unknown whether distinct root architectures appear when the intercropping plants respond to different P levels. In the present study, we used a custom apparatus consisting of a transparent manual root box to observe the root system in situ in a non-destructive manner. We tested how root distributions and root tendencies changed in tomato/potato onion intercropping systems with no P added or 120 mg·kg^−1^ P added. We measured changes in tomato plant growth and analyzed how plant interactions affected cropping patterns and P levels. We hypothesized that the spatial root distribution of the tomato and potato onion plants would be compatible yet distinct at both P levels, contributing to increased plant growth.

## RESULTS

### Influence of cropping patterns and P levels on tomato plant growth

The cropping pattern and P level treatments both affected shoot biomass and total plant biomass in the tomato plants, and we observed a significant P level × cropping pattern interaction for shoot biomass and plant total biomass (P>0.05; Table 1). Compared with the biomass of monoculture tomato plants, the shoot biomass and total plant biomass of the tomato plants increased significantly when intercropped with potato onion for both no P added and 120 mg·kg^−1^ P added treatments. For tomato root biomass, we did not detect any statistically significant interactions between the cropping pattern and P level treatments. However, tomato root biomass was clearly influenced by the cropping pattern (P>0.05; Table 1), although it was not significantly influenced by the P level.

**Table 1.** Effect of intercropping with potato onion on tomato’s plant growth (mg plant ^−1^) at different P levels.

|  | P0 |  | P120 |  | F-statistics |  |  |
| --- | --- | --- | --- | --- | --- | --- | --- |
|  | Monocultu<br>re | Mixed<br>culture | Monocultu<br>re | Mixed<br>culture | P level | Cropping<br>pattern | P levelx<br>Cropping<br>pattern |
| Shoot dry<br>weight (mg<br>shoot<sup>-1</sup>) | 0.73±0.05<br>b | 0.99±0.02<br>a | 1.20±0.08<br>b | 1.65±0.07<br>a | **263.83\*\*\*** | **104.56\*\*\*** | **7.61\*** |
| Root dry<br>weight (mg<br>root<sup>-1</sup>) | 0.18±0.02<br>a | 0.21±0.01<br>a | 0.17±0.01<br>a | 0.20±0.03<br>a | 0.39 | **7.47\*** | 0.09 |
| Total plant<br>dry weight<br>(mg plant<sup>-1</sup>) | 0.91±0.04<br>b | 1.20±0.02<br>a | 1.37±0.08<br>b | 1.85±0.09<br>a | **214.12\*\*\*** | **102.80\*\*\*** | **5.89\*** |

P0 represents no P added treatment, P120 represents 120 mg·kg<sup>-1</sup> P added treatment. \*, \*\*, \*\*\* represent sterilization contrasts significant at the 0.05, 0.01, and 0.001 probability levels, respectively.

### Root length density distribution

Influenced by both neighboring plants and the P application rate, RLD was unevenly distributed in the mixed cultures (Fig. 1A, B, G, and H) but evenly distributed in the tomato monoculture (Fig. 1E and K) and the potato onion monoculture (Fig. 1C and L). When tomato plants were intercropped with potato onion plants, the RLD area of tomato (Fig. 1A and G) was much deeper than in the tomato monoculture (Fig. 1E and K). In the mixed cultures, the roots of the tomato plants (Fig. 1A and G) always avoided contact with the roots of the potato onions, whereas the roots of the potato onion (Fig. 1B and H) spread laterally under the neighboring tomato plants under both P application treatments.

**Fig. 1.**
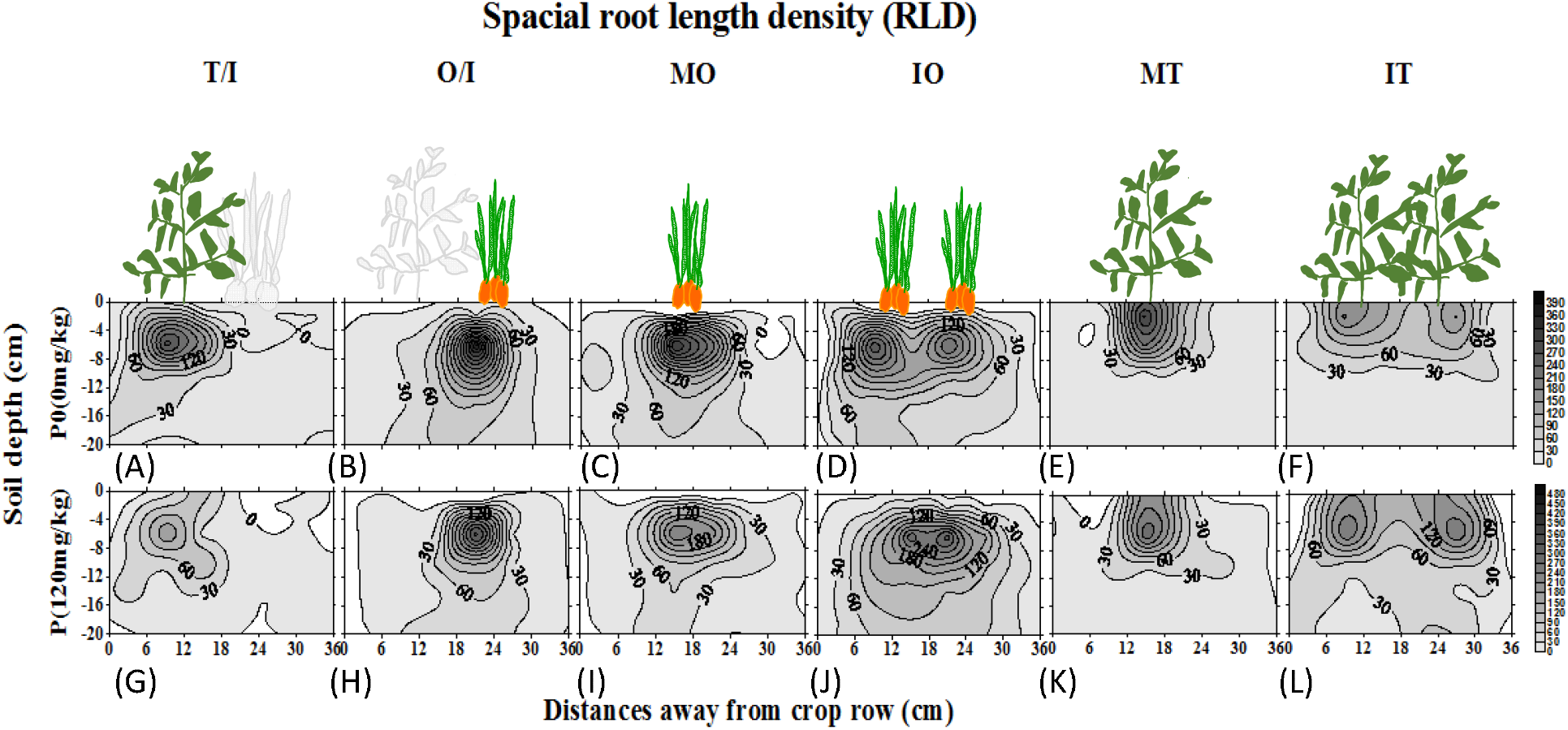
Spatial root length density (RLD) (cm per 120 cm^3^ soil volume) for various treatments [(A, G) intercropped tomato, (B, H) intercropped potato onion, (C, I) one potato onion plant monoculture, (D, J) two potato onion plants monoculture, (E, K) one tomato plant monoculture, (F, L) two tomato plants monoculture] under no P added and 120 mg·kg^−1^ P added treatments. The contour lines are at intervals of 1 cm/12 cm^3^ soil volume.

The root tendency of plants neighboring the same species was more strongly influenced by P level. In the no P added treatment, the RLD areas of the potato onion and tomato were not intermingled when neighboring the same plant species (Fig. 1D and F). By contrast, for the 120 mg·kg^−1^ P added treatment, the RLD areas of potato onion plants neighboring the same species (Fig. 1J) were markedly intermingled, whereas the RLD areas of tomato plants neighboring the same species (Fig. 1L) were clearly separated.

### Root weight density distribution

The root weight density distribution is shown in Fig. 2. Under the no P added and 120 mg·kg^−1^ P added treatments, the 0.2 g kg^−1^ soil root weight density (RWD) contour of the intercropped tomato plants (Fig. 2A and G) occupied a deeper soil layer than that of the tomato monoculture, and under the no P added treatment, the RWD of the tomato plants was higher than in the monoculture. The 2 g/kg soil RWD contour distribution of potato onion plants (Fig. 2B and H) was distributed in a narrower area than in the monoculture (Fig. 2C and I), and the RWD of the intercropped potato onion plants (Fig. 2B and H) was higher than for the monocultures (Fig. 2C and I). The directions of RWD in both intercropping systems and monocultures were the same as for RLD.

**Fig. 2.**
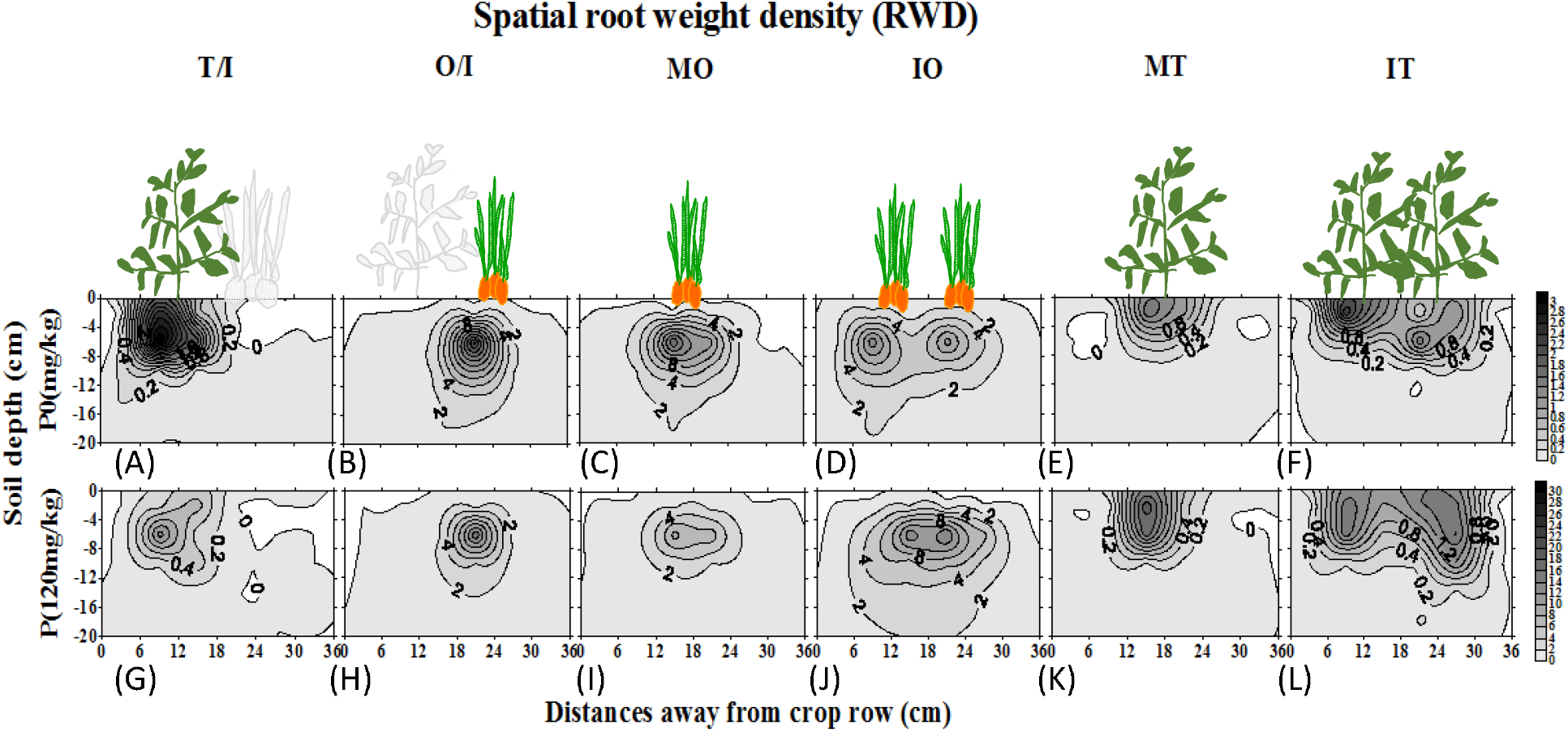
Spatial RWD (g/kg) for various treatments [(A, G) intercropped tomato, (B, H) intercropped potato onion, (C, I) one potato onion plant monoculture, (D, J) two potato onion plants monoculture, (E, K) one tomato plant monoculture, (F, L) two tomato plants monoculture] under no P added and 120 mg·kg^−1^ P added treatments. The contour lines are at intervals of 1 g root fresh weight per kilogram fresh soil.

In general, the root distributions of the tomato plants became deeper when intercropped with potato onion plants than those under tomato monoculture, and under the no P added conditions, the RWD of the tomato plants became higher than in monoculture. For the two P levels, the root tendencies of the two crops were significantly different, with the tomato roots avoiding contact with potato onion roots and the potato onion roots clearly extending towards the tomato roots. When the tomato plants were next to the same species, their roots were crossed under the no P added treatment, whereas they avoided crossing under the 120 mg·kg^−1^ P added treatment. The root distribution of the potato onion plants in the intercropping system became narrower than in monoculture. Additionally, the root tendency of the potato onion plants neighboring the same species was opposite to that of the tomato roots: under the no P added treatment, the root areas of the potato onion plants neighboring the same species were separated, whereas under the 120 mg·kg^−1^ P added treatment, the roots were significantly crossed.

### Root tendency in root boxes

Image data were obtained on the 12th day after transplantation (sampling time was tested in our previous experiment to ensure that the roots of two plants in one box remained uncrossed). Fig. 3 shows how the root architecture was affected by neighboring plants under the no P added treatment in the root box. In P0MT1 (one tomato plant in monoculture) and P0MO1 (one potato onion plant in monoculture), the roots of tomato plants and potato onion plants were distributed evenly. In the other combinations, the roots were unevenly distributed to a significant extent. In P0I (the tomato/potato onion intercropping system), the tomato roots avoided the potato onion roots significantly, whereas the potato onion roots spread laterally under the tomato row. In P0MT2 (two tomato plants in monoculture), the roots of the two tomato plants were clearly crossed, whereas the potato onion roots avoided other potato onion roots in P0MO2 (two potato onion plants in monoculture).

**Fig. 3.**
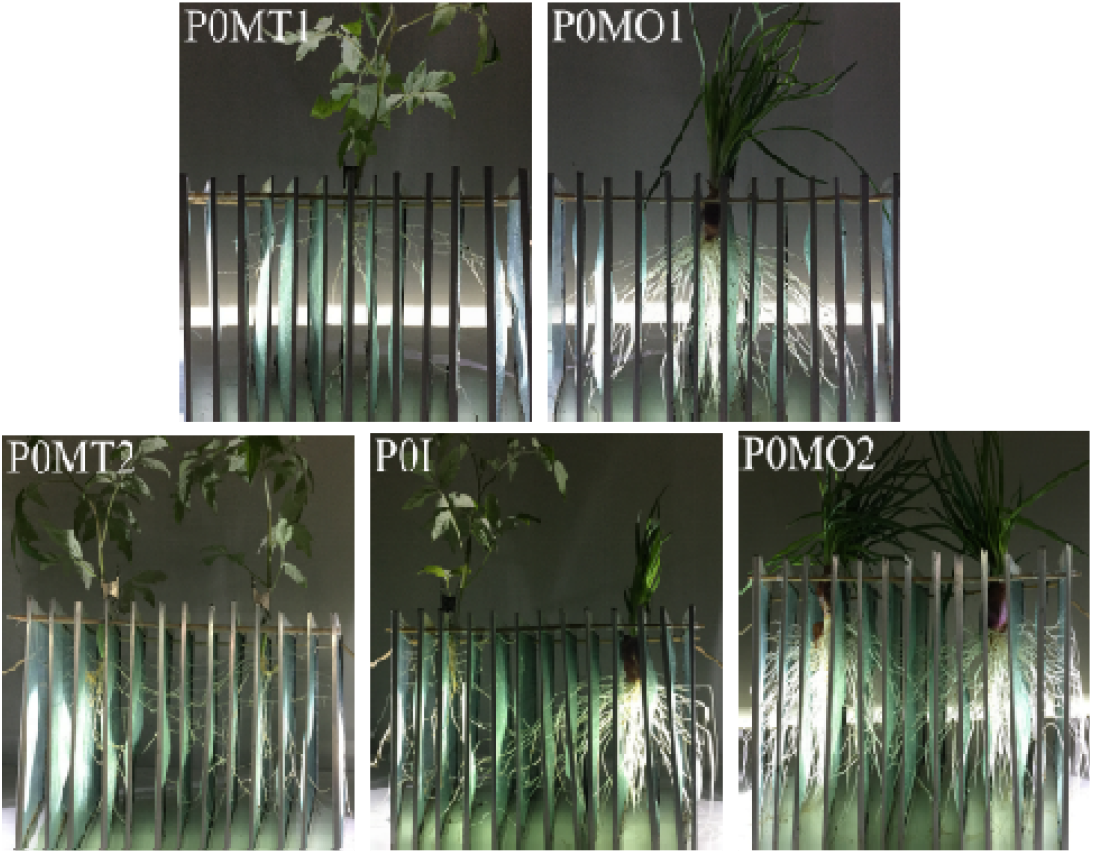
Images of roots from different tomato/potato onion combinations under no P added conditions at day 12. (P0MT1) one tomato plant in monoculture, (P0MO1) one potato onion plant in monoculture, (P0MT2) two tomato plants in monoculture, (P0I) tomato/potato onion intercropping system, (P0MO2) two potato onion plants in monoculture.

For the 120 mg·kg^−1^ P added treatments (Fig. 4), the root distributions for the different treatment combinations were different from those under the no P added treatments. In PMT1 (one tomato plant in monoculture) and PMO1 (one potato onion plant in monoculture), the tomato and potato onion roots were evenly distributed, which is associated with P deficiency. In PI (the tomato/potato onion intercropping system), the tomato roots also avoided the potato onion roots, whereas the potato onion roots spread in a nearly uniform distribution. In PMT2 (two tomato plants in monoculture) and PMO2 (two potato onion plants in monoculture), the root tendencies of the tomato plants and potato onion plants exhibited opposite trends to those under the no P added treatment; in PMT2, the tomato roots avoided intermingling, whereas the potato onion roots showed no significant trend.

**Fig. 4.**
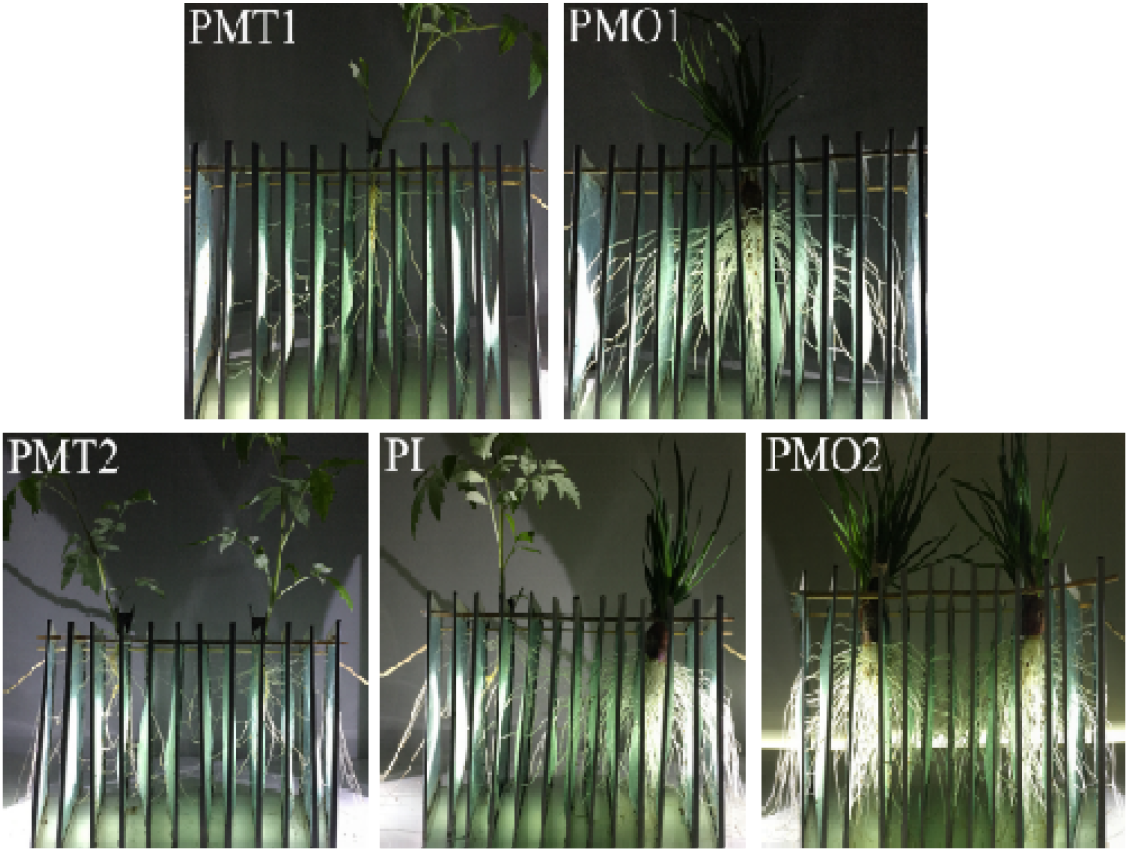
Images of roots from different tomato/potato onion combinations under 120 mg·kg^−1^ P added conditions at day 12. (PMT1) one tomato plant in monoculture, (PMO1) one potato onion plant in monoculture, (PMT2), two tomato plants in monoculture, (Zhang et al.) tomato/potato onion intercropping system, (PMO2) two potato onion plants in monoculture.

### Root percentage distribution in root boxes

Consistent with Figs. 3 and 4, the distribution of the root percentage is shown in Fig. 5. In the tomato/potato onion mixed culture, the tomato root length percentage of P0IT in space 2 was significantly higher than in PIT, and the tomato root length percentage of P0IT in the 6–9-cm spaces was lower than in PIT, indicating that tomato roots avoided potato onion roots more clearly than under the no P added treatment. In the P0MT2 treatment, the root length percentage in the middle area was higher than on the two sides, with horizontal distance 16-cm showing the highest percentage. In PMT2, the root length percentage was higher in horizontal distance 2–6-cm and 24–28-cm than in 12–18-cm. Specifically, when tomato plants were intercropped with the same species, their root growth trend was related to the P level, with the roots crossing under the no P added treatment, whereas they clearly avoided one another under the 120 mg·kg^−1^ P added treatment.

**Fig. 5.**
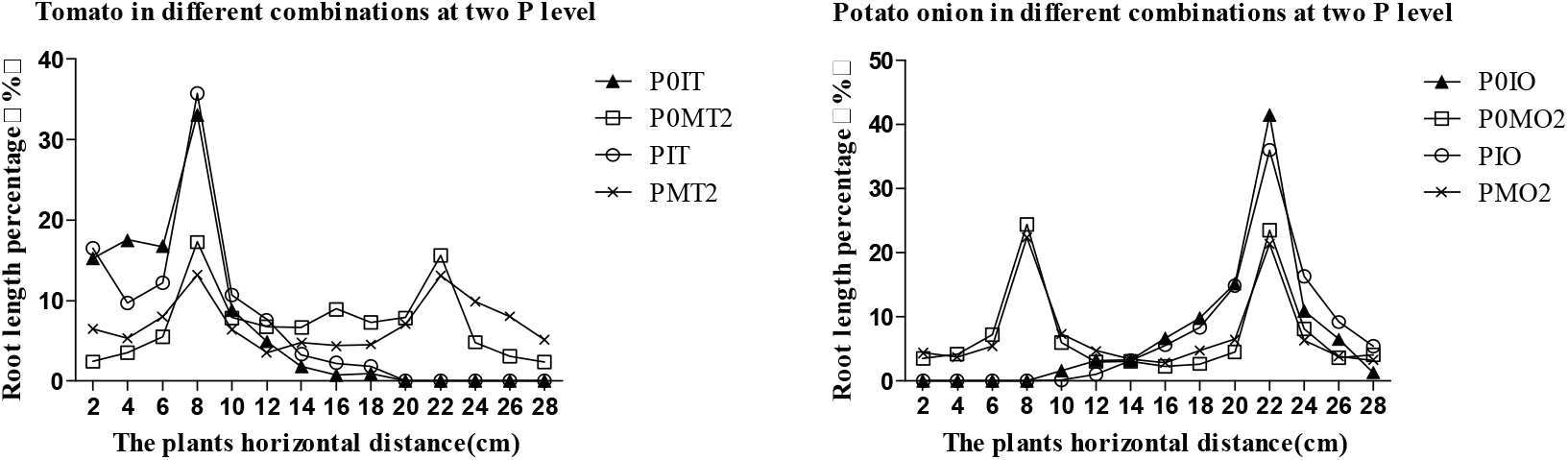
Root percentage data for tomato and potato onion grown in different combinations in 14 spaces. The distribution of the tomato root percentage is shown on the left and that of the potato onion root percentage is shown on the right. (P0IT) tomato in the intercropping system under no P added, (P0MT2) tomato monoculture under no P added, (PIT) tomato in the intercropping system under 120 mg·kg^−1^ P added, (PMT2) tomato monoculture under 120 mg·kg^−1^ P added; (P0IO) potato onion in the intercropping system under no P added, (P0MO2) potato onion monoculture under no P added, (PIO) potato onion the in intercropping system under 120 mg·kg^−1^ P added, (PMO2) potato onion monoculture under 120 mg·kg^−1^ P added.

Fig. 5 shows that the root length percentage of potato onion plants in P0IO was higher on the left than on the right, and the root length percentage in horizontal distance 10–18-cm was higher than in PIO. In PIO, the potato onion roots were distributed evenly on both sides, revealing that the potato onion roots tended to extend towards the tomato roots when the species were intercropped under the no P added treatment, although this trend was not significant under the 120 mg·kg^−1^ P added treatment. In P0MO2, the root length percentages on both sides (2–4-cm and 26–28-cm) were higher than in the middle zone (12–18-cm), whereas in PMO2, there was no obvious trend. In other words, when potato onion plants were intercropped with the same species under no P added treatments, the roots avoided mixing, whereas there was a less obvious trend under the 120 mg·kg^−1^ P added treatments. All of these data are consistent with the results presented in Figs. 3 and 4.

## DISCUSSION

Enhancement of plant biomass under intercropping has been observed in many experiments (Awal et al., 2006; Li et al., 1999, 2001; Tsubo and Walker, 2002; Xia et al., 2013; Zhang et al., 2007). In our experiments, results for both P treatments showed that aboveground tomato biomass and plant total biomass could both be increased significantly by intercropping with potato onion plants. We detected statistically significant interactions between the cropping patterns and P level treatments for both shoot biomass and total plant biomass of tomato in the intercropping system. Regarding the root dry weights of the tomato plants, we did not detect any statistically significant interactions between cropping pattern and P level treatments, although this metric was clearly influenced by cropping pattern (P>0.05) and not significantly influenced by the P level. In previous studies, few experiments have addressed the effects of the interaction between cropping patterns and P levels on plant growth in intercropping systems. However, Li et al. (2008) and Wang et al. (2007) have shown that root biomass can be differentially affected in different intercropping system combinations, increasing in some contexts while remaining the same as in monocultures in others, suggesting that in mixed cultures, root biomass can be influenced by the species type of adjacent plants. In addition, Li et al. (2010) showed that the root biomasses of different crops can vary under different P levels. Thus, the root biomass in intercropping systems can be affected by many factors. In our experiment, the root biomass of tomato plants was significantly influenced by intercropping tomato plants with potato onion plants.

Root interactions have been studied in many intercropping systems, and the spatial distribution of roots and their density in the soil has been shown to determine the ability of a crop to acquire the necessary nutrients and water to sustain growth. In this study, the roots of tomato plants and potato onion plants both showed an extended root distribution, and the RLD and RWD of the tomato plants both became deeper than in tomato monoculture, consistent with previous literature (Adiku et al., 2001; Gao et al., 2010; Li et al., 2006). Some studies have found that the root distribution can become unbalanced and that roots can extend horizontally to greater distances in an intercropping system (Zhang and Huang, 2003), with overyielding of species resulting from the greater lateral deployment of roots and increased RLD. The roots of intercropped plants can extend into the root area of other plants and sometimes penetrate deeper than in monoculture (Adiku et al., 2001; Li et al., 2006), and the compatibility of the spatial root distributions of the intercropped species contributes to interspecific facilitation. In our experiment, the extension of the root distribution and the deeper root space of tomato plants may have contributed to increased plant biomass. However, the root tendencies of the two crops observed here were not the same as in previous studies. In previous studies by Adiku et al. (2001) and Li et al. (2006), the roots of two crops were found to extend into each other’s root areas. Thus, we believe that the root tendency of one plant in an intercropping system may be influenced by both plant species, and whether the roots of the two crops are mixed or separate, these different tendencies are beneficial for plant nutrient absorption. These tendencies generally involve an extended root distribution, and greater root length can help plants absorb nutrients and increase biomass.

In previously studies, the results of competition have always been connected to resources and plant species. Some authors believe that the results of competition can be variable in different environments, with intraspecific competition being dominant under some conditions (Sheley and Larry, 1994; Velagala et al., 1997; Wassmuth et al., 2009), whereas interspecific competition is stronger under others (Blank, 2010; Vasquez et al., 2008; Young and Mangold, 2008). Ge et al. (2000) demonstrated that low inter-root competition is a more efficient way for adjacent plants to decrease root overlap, and Zhang et al. (2002) showed that when root weight is at its maximum and roots do not overlap in a wheat/faba bean intercropping stage, then competition between the two crops for water and nutrients can be reduced, resulting in higher yields for both species. In our intercropping system, the root action of tomato plants was consistent with that observed for this previous study, and under a nutrient-deficient conditions, the roots opted to decrease their overlap and decrease their competition with potato onion plants.

In our analysis of root tendency, when the tomato and potato onion plants were planted with their same species, the reaction of the roots was more closely related to the P level. The roots of the potato onion plants were clearly separated from those of their same-species neighbors under no P added treatment, which appeared to aid in avoiding competition and improving survival of the species, whereas no obvious root tendency was observed in the absence of P stress. However, the tomato roots intermingled with one another under the no P added treatment, possibly allowing them to compete for more resources, whereas the roots clearly avoided intermingling under the 120 mg·kg^−1^ P added treatment, possibly to avoid competing for resources. Cheplick and Kane (2004) reported when two kin plants are planted together, their roots can avoid one another or engage in spatial segregation to avoid competing for resources, whereas when non-kin plants are planted together, the roots usually overlap, allowing for more competition. In these experiments, the root behavior of the potato onions was consistent with previous results, perhaps because in this variety was not subjected to artificial transformation. Or in other words, for the potato onion, the results regarding root recognition appeared biased towards protecting the species itself, thus preventing competition among roots under no P added treatment. However, the responses of the tomatoes were different from those of Cheplick and Kan. Generally, studies on kin recognition have been conducted on wild plants, whereas few such studies have been performed on crop species (Dudley and File, 2007; Murphy and Dudley, 2009). Wild plants usually grow under natural conditions in which resources are limited; however, in some long-term cultivated species grown under resource-rich conditions (Wenke, 1980), the ability of roots to recognize those of their kin have gradually decreased, and root recognition can be affected by plant species and genotype in a significant manner (Fang et al., 2011). Therefore, considering that the tomato seeds we selected have been subjected to long-term cultivation, we speculate that the root recognition may have been weakened in these plants. Or in other words, when planted with their siblings under nutrient-deficient conditions, these plants no longer know to protect their kin. The results of our study clearly show that tomato and potato onion roots can respond to nutrients and adjacent plants, consistent with the viewpoint of Cahill et al. (2010), although determining which factors in an intercropping system are most important for controlling root behavior requires further research.

## MATERIALS AND METHODS

### Plant materials and cultivation conditions

The tomato (*Solanum lycopersicum L*.) variety “Dongnong708” was provided by the Tomato Breeding Center of Northeast Agricultural University (Harbin, China). The potato onion (*Allium cepa* var. *agrogatum Don*.) variety Suihua, a native variety with potential allelopathy (Liu et al., 2013), was provided by the Laboratory of Vegetable Physiological Ecology (Harbin, China). Tomato seeds were treated with hot (55°C) water and germinated in Petri dishes with wet gauze in the dark at 28°C. Seedlings with two cotyledons were planted in plastic pots (8×8 cm) containing 100 g soil after emergence, and seedlings with four leaves were then used for the different experiments. All of the seedlings were cultivated in a phytotron located in the Experimental Center at Northeast Agricultural University, Harbin, China (45°41′N, 126°37′E), from July 2014 to October 2015, and the phytotron was maintained under the following conditions: 14/10 h light/dark cycle, 28/18°C day/night temperature and 70% air relative humidity. The potato onion plants were stored at 4°C before planting. All of the experiments were performed in the Laboratory of Vegetable Physiological Ecology (Harbin, China).

### Experiment 1: Tomato/potato onion mixture in pots

The primary pot treatments consisted of no additional added P and 120 mg·kg^−1^ P added. These P concentrations were based on previous experimentation showing that soil with no additional P is insufficient for tomato growth and that soil with 120 mg·kg^−1^ P added is sufficient. The sub-pot treatments addressed tomato/potato onion intercropping and tomato monoculture in plastic pots (28 cm diameter, 20 cm height) containing 3 kg soil. At the time of tomato transplantation, the potato onion plants were planted, and the tomato:potato onion ratio was 1:3 in the intercropping treatment. The experimental design was a randomized complete block design with three replicates. Four treatments were performed in each block, and 4 pots were included in each treatment. In all, there were 16 pots per block and with 3 blocks total, yielding 48 pots in all. Each pot was watered with tap water every 3 days to maintain the soil water content at approximately 60% of the water-holding capacity, and the plants were grown in the phytotron as described above.

Sandy loam soil from the 30–50 cm layer under the ground surface was collected from an open field at Northeast Agricultural University (Harbin, China). The soil contained 17.4 g·kg^−1^ organic matter, 40.6 mg·kg^−1^ available N (nitrate and ammonium), 11.4 mg·kg^−1^ Olsen P and 100.9 mg·kg^−1^ available K, and it exhibited an electrolytic conductivity (1:5, w:v) of 153.5 mS·cm^−1^ and a pH (1:5, w:v) of 6.98. Previous experiments have shown that even when the total P and available P levels are relatively high, soil can still be considered P-deficient for plants if plant growth can be improved by P addition (Holloway et al., 2001; Li et al., 2005; Wang et al., 2007). In a previous experiment, we confirmed that the base soil P content was insufficient for tomato plants (data not shown).

P was added as KH_2_PO_4_ at 120 ppm for the 120 mg·kg^−1^ P added treatment, and fertilization with 120 ppm N (in the form of CO(NH2)2) and 120 ppm K (in the form of K_2_SO_4_) was performed for both the no P added and 120 mg·kg^−1^ P added treatments; then K_2_SO_4_ was used to balance the K rate for the two P level treatments. Plants were harvested on the 20th day after transplantation, thoroughly washed with distilled water and separated into roots and shoots. The shoots and roots were killed by heating at 105°C for 30 min and then dried at 60°C for 72 h.

### Experiment 2: Tomato/potato onion mixture in foam boxes

The same soil and fertilizer management techniques described above were used in this experiment. To provide sufficient space for the plant roots and to reduce harm to the root system when sampling, we employed large foam boxes with an internal volume of 36×25×22 cm as culture pots. Each foam box was filled with 20 kg soil. The experimental design was a randomized complete block design with two replicates, and ten treatments were used in this experiment. The primary pot treatments were no P or 120 mg·kg^−1^ P added, and the sub-pot treatment consisted of five intercropping combinations: 1) a tomato/potato onion intercropping system, 2) one tomato plant in monoculture, 3) two tomato plants in monoculture, 4) one potato onion in monoculture and 5) two potato onion plants in monoculture. The tomato to potato onion ratio in the intercropping treatment was 1:3. When considering the nutrient balance per each box, the three potato onion plants were viewed as equivalent to one plant.

The plants were sampled on the 20th day after transplantation, and root samples were collected using the monolith method, as modified by Li et al. (2006) and Smit et al. (2013). Briefly, the foam box was cut into vertical sections at 10-cm intervals along the wide side, the soil surface was made as smooth as possible, and the roots were then fixed in each 6×4-cm area with 5 cm nails. Finally, a 6×5×4 cm inner-diameter iron box was used to remove a 5 cm layer of soil from the center of the foam box; the volume of each soil block was 120 cm^3^. There were 30 monoliths (5 in a vertical and 6 in a horizontal direction) in each soil profile, and 600 monoliths were sampled in total. Each soil sample was placed in a numbered plastic bag.

All of the soil samples were poured onto a sieve (0.2 mm mesh, 30 cm diameter, 5 cm height) and stirred until all of the roots could be freed of soil using very fine tweezers. The sieves were suspended in a large water bath and shaken continuously, and the soil material remaining in the sieves was removed manually. The tomato and potato onion roots were distinguished by differences in color, smell and fibrous roots. For example, tomato roots are yellowish and hairy, whereas potato onion roots have a smooth surface with white coloration and some degree of transparency.

### Experiment 3: Tomato/potato onion mixture for imaging the root tendency in the root boxes

This experimental design was the same as experiment 2, and the treatments were assessed in a transparent manual root box (a practical invention patent application has been filed for this box). The root box consisted of transparent glass pieces and thirteen layers of nylon mesh with a 1 mm aperture, allowing us to image root architecture in situ without destroying the roots. Each root box was filled with a 15 kg mixture consisting of one part sand and 3 parts vermiculite. For the experiment, the plants were irrigated with modified Hoagland's nutrient solution, and the no P added and 120 mg·kg^−1^ P added treatments were created by adding P_2_O_5_ and KH_2_PO_4_ at 80 μM and 320 μM concentrations, respectively. N and K were applied as CO(NH_2_)_2_ and K_2_SO_4_, respectively, at 100 ppm, and K_2_SO_4_ was used to balance the K rate in both treatments. Other nutrients were provided at the concentrations indicated by Fontes et al. (1986).

At the time of sampling, we removed the bottom of the root boxes and soaked them in water. Half an hour later, when the sand and vermiculite were almost washed free, the root box was gently removed and the culture medium was thoroughly rinsed from the root surfaces by spraying. Two wooden sticks were run through the 13 networks from two sides, and the wire frame was fixed with plastic grips. Two flashlights were used as a light source when taking photographs. After imaging, the roots between every two grids were cut with a pair of scissors as an individual sample, and the root length of each sample was determined using a root system scanner, which we used to calculate the root percentage.

### Statistical analysis

Results regarding plant growth were analyzed using the SAS 8.0 software program (SAS Institute Inc., Cary, USA), and the means of the different treatments were compared using Tukey’s test at the *p* = 0.05 level. The data are expressed as the means with standard errors. We used general linear models to determine the significance of the primary effects (P level and cropping pattern) and interactions (P level × cropping pattern) on tomato plant growth. Data from the monoliths from the white foam box experiment represent the entire root population in each soil profile. The results are presented as contour diagrams. Root length density (RLD) contour diagrams were prepared using the Surfer v. 8.0 software program (Golden Software Inc., Golden, CO). Images from the root box experiment were obtained in panoramic view using an Apple mobile phone. The root length percentage was determined using an image scanner analyzer (LA - S2400).

## CONCLUSIONS

Our study provides novel findings regarding plant growth and root interactions in an intercropping system under differing P levels. Based on our results, we conclude that deeper and more evasive root distributions in tomato plants can support greater tomato biomass in a tomato/potato onion intercropping system.

## ACKNOWLEDGEMENTS

We would like to thank Dr. Long Li (Department of Plant Nutrition, China Agricultural University) for his valuable guidance in our experiment’s design and result analysis.

## SYMBOLS AND ABBREVIATIONS

Phosphorus-P

